# Integrative genome-scale analyses reveal post-transcriptional signatures of early human small intestinal development in a directed differentiation organoid model

**DOI:** 10.1101/2022.07.12.499825

**Authors:** Yu-Han Hung, Meghan Capeling, Jonathan W. Villanueva, Matt Kanke, Michael T. Shanahan, Sha Huang, Rebecca L. Cubitt, Vera D. Rinaldi, John C. Schimenti, Jason R. Spence, Praveen Sethupathy

## Abstract

MicroRNAs (miRNAs) are important post-transcriptional gene regulators in organ development. To explore candidate roles for miRNAs in prenatal SI lineage specification in humans, we used a multi-omic analysis strategy in a directed differentiation model that programs human pluripotent stem cells toward the SI lineage. We leveraged small RNA-seq to define the changing miRNA landscape, and integrated chromatin run-on sequencing (ChRO-seq) and RNA-seq to define genes subject to significant post-transcriptional regulation across the different stages of differentiation. Our analyses showed that the elevation of miR-182 and reduction of miR-375 are key events during SI lineage specification. We demonstrated that loss of miR-182 leads to an increase in the foregut marker *SOX2*. We also used single-cell analyses in murine adult intestinal crypts to support a life-long role for miR-375 in the regulation of *Zfp36l2*. Finally, we uncovered opposing roles of SMAD4 and WNT signaling in regulating miR-375 expression during SI lineage specification. Beyond the mechanisms highlighted in this study, we also present a web-based application for exploration of post-transcriptional regulation and miRNA-mediated control in the context of early human SI development.

**Graphical Abstract:** 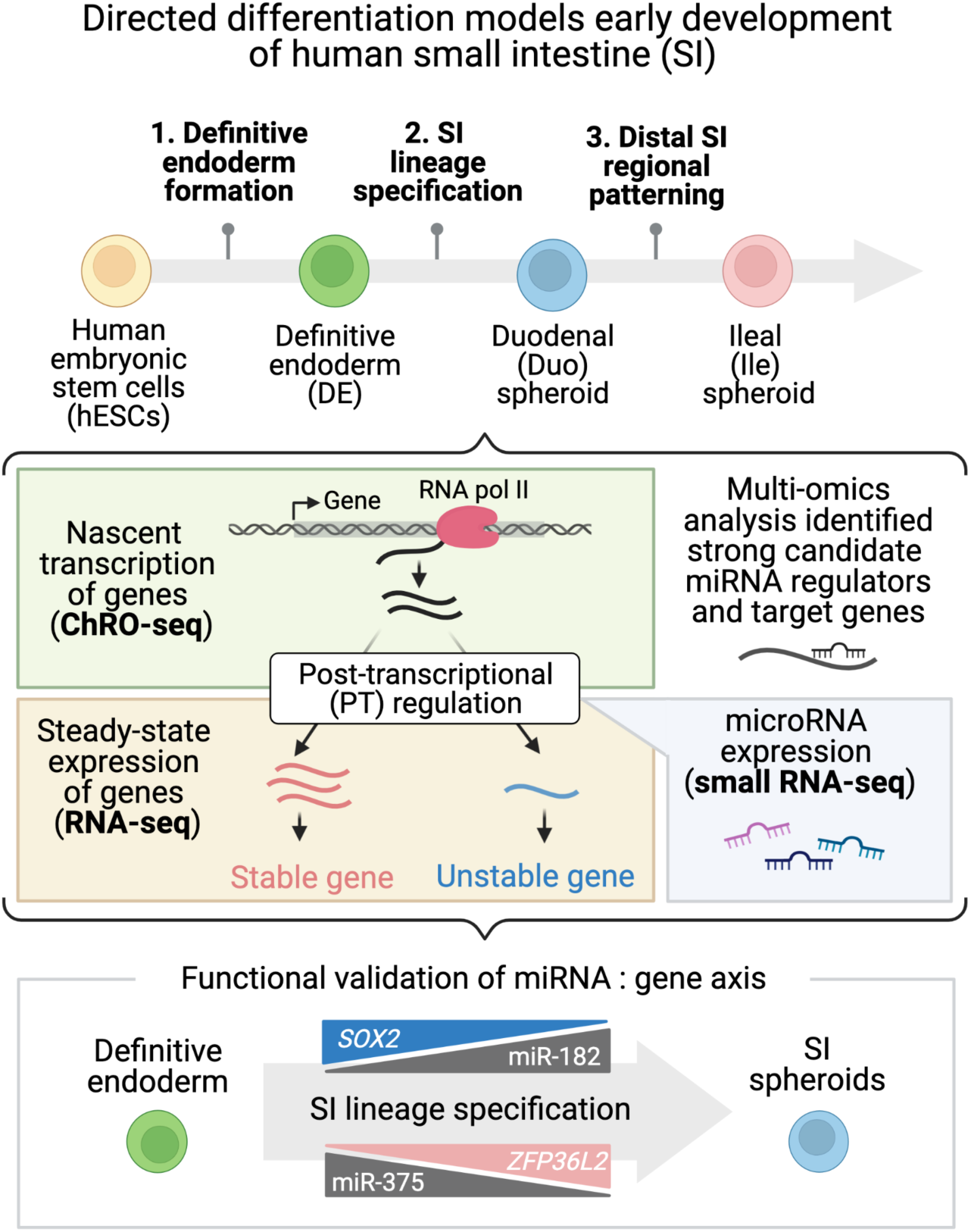

## Introduction

MicroRNAs (miRNAs) are small, non-coding RNA molecules (∼22 nucleotides) that function as post-transcriptional gene suppressors in various biological processes including development (1, 2). The biogenesis of mature miRNAs occurs through a multi-step process. Longer primary miRNA transcripts are usually transcribed by RNA polymerase II and subsequently processed by enzymes into mature, functional miRNAs. Mature miRNAs confer gene silencing through complementary base pairing with mRNA target sites (typically in 3′ untranslated region of transcripts), leading to destabilization and degradation of the target mRNAs (3). As demonstrated by several seminal studies, miRNAs play an essential role in controlling dynamics of gene expression during mammalian development. Murine embryonic lethality was observed when disrupting miRNA maturation (4, 5) or function (6) *in vivo*. In addition, different fetal tissues were shown to have distinct miRNA signatures and temporal dynamics in both mouse and humans (7), suggesting that miRNAs participate in lineage specification during pre-natal development.

This study sought to identify candidate miRNA regulators of early small intestinal (SI) development in humans. While specific miRNAs have been linked to the regulation of embryonic development of various organs such as the heart (8-11), brain (12-15), pancreatic islets (16-18) and skeletal muscle (19-21), the role of miRNAs in SI development has not been as closely studied. Mice with intestinal epithelial-specific knockout of Dicer (a key miRNA processing enzyme) were reported to exhibit disrupted gut epithelial architecture and function (11), conveying the importance of miRNAs to proper SI development. However, the roles of specific miRNAs in the context of SI development are unexplored in humans. Directed differentiation from human pluripotent stem cells (hPSCs) to human intestinal organoids (HIOs) represents a state-of-art model to study human SI development (22-24). To generate HIOs, hPSCs are differentiated into definitive endoderm, which results in the formation of 3-dimensional (3D) spheroids fated to the SI lineage, which then develop into more mature HIOs. These SI spheroids can be regionally patterned (duodenum vs. ileum) by controlling the length of time that they are exposed to FGF and WNT signaling (23). We have previously applied chromatin run-on sequencing (ChRO-seq)(25) in this model system to successfully profile enhancer dynamics and define transcriptional programs relevant to SI developmental events in human (26). Here we aimed to define post-transcriptional programs that underlie human SI development and identify candidate master miRNA regulators in this context.

To that end, we developed a multi-omic analysis strategy that integrates ChRO-seq, RNA-seq and small RNA-seq (smRNA-seq) generated from the hPSC-HIO model. Specifically, we first used smRNA-seq to characterize the dynamics of the miRNA landscape across different cellular stage transitions of SI differentiation. Next, we integrated ChRO-seq (which measures nascent gene transcription) and RNA-seq (which measures steady-state gene expression) to tease apart the contribution of transcriptional and post-transcriptional (PT) regulation for each gene individually. We made quantitative measurements of PT regulation and then identified the miRNAs that likely mediate the most PT regulation during endoderm formation, SI lineage specification, and SI regional patterning. Our analyses pointed to the gain of miR-182 (and the consequent loss of *SOX2*) and the loss of miR-375 (and the consequent gain of *ARL4C* and *ZFP36L2*) as key events during SI lineage specification. Further analyses showed that whereas miR-182 levels are likely controlled by PT mechanisms, miR-375 is regulated robustly by transcription during human SI lineage specification. Follow-up functional studies in the hPSC-HIO model confirmed that miR-182 suppresses the foregut driver gene *SOX2*. In addition, single cell RNA-seq of adult miR-375 knockout mice supported a continued, post-developmental role of miR-375 in the regulation of *ARL4C* and *ZFP36L2*. In this study, by using an integrative multi-omics approach, we offer the first look into PT regulation during early human SI development and provide novel insights into the function of miRNAs in this context. We also provide the research community with a searchable web server (https://sethupathy-lab.shinyapps.io/ProHumID/) to explore the omics data we generated from the hPSC-HIO model.

## Results

### SmRNA-seq shows a changing miRNA landscape during human SI lineage commitment

To study early SI development in humans, we carried out directed differentiation from human embryonic stem cells (hESCs) toward the SI lineage as previously described (22, 23, 27)(**Figure 1A**). Cells from the stage of hESC, definitive endoderm (DE), duodenal (Duo; proximal SI) spheroids, and ileal (Ile; distal SI) spheroids were collected to study molecular changes (**Figure 1A**). To characterize the dynamics of miRNA expression, we performed smRNA-seq across these stages. Principal component analysis (PCA) and unsupervised hierarchical clustering demonstrated distinct miRNA profiles at each stage (**Figure 1B and C**). We observed that miRNAs (RPMMM > 1000 in at least one stage) exhibit changing expression patterns over the course of directed differentiation (**Figure 1D**). Consistent with previous microarray and smRNA-seq studies comparing hESC and DE (28-30), we found several expected miRNAs enriched in hESCs (e.g., miR-512b, miR-222, miR-7) or in DE (e.g., miR-708, miR-375, miR-489, miR-9)(**Figure 1D**). In addition, previous reports showed that the miR-302/367 family is critical for maintaining stem cell pluripotency (31-33) and that miR-302 isoforms may have functional roles during differentiation of hPSCs to DE (28). Our sequencing data indeed confirmed enrichment of miR-302 family members and isoforms in the hESC and/or DE stages (**Figure 1D**).

**Figure 1.**
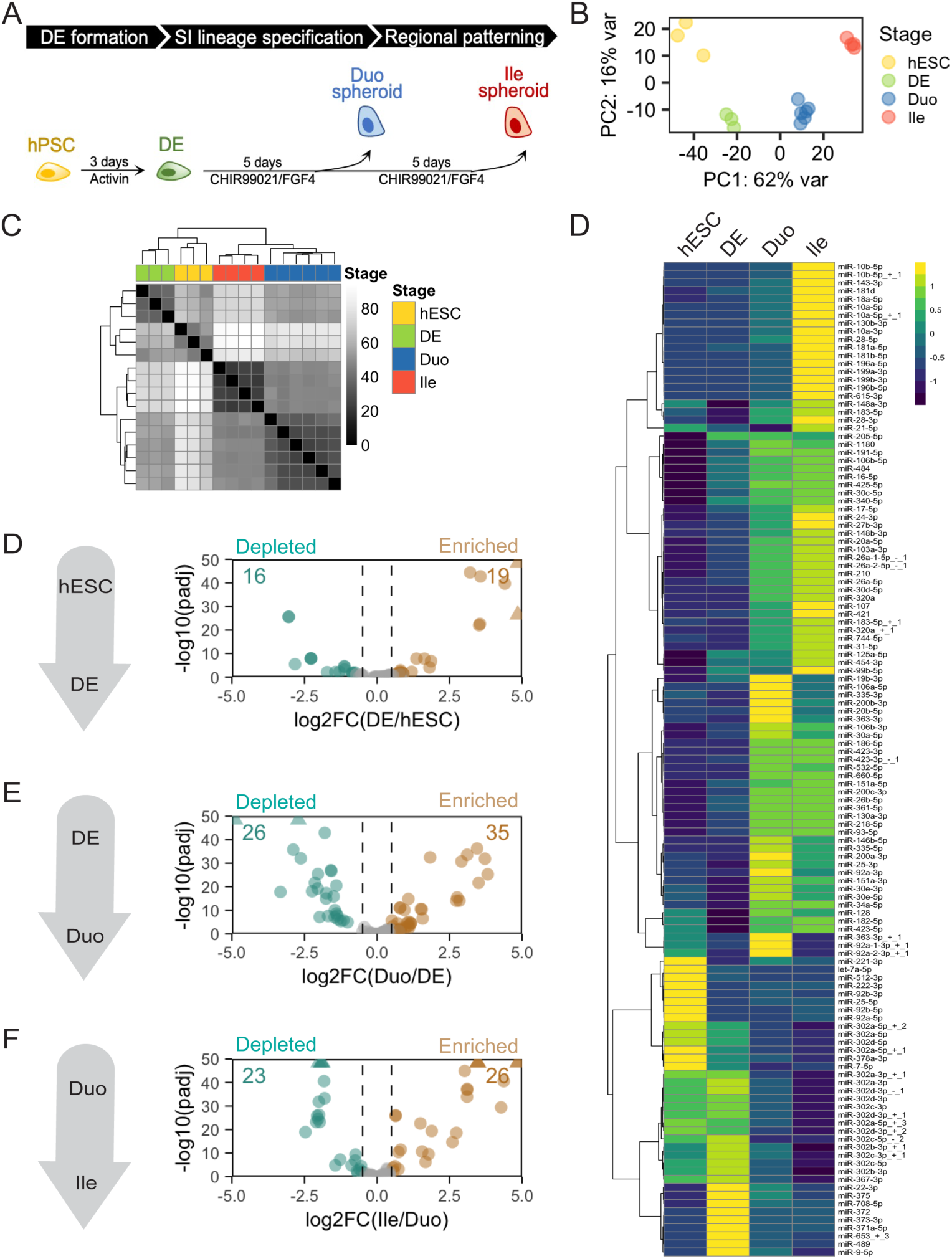
smRNA-seq revealed changing miRNA signatures during early SI development in a human organoid model. (A) The stage transitions of the hPSC-HIO model recapitulate early events of human SI development: DE formation, SI lineage formation and SI regional patterning. (B) PCA of smRNA-seq miRNA profiles of hESC, DE, Duo and Ile spheroids. (C) Hierarchical clustering analysis (Euclidean distance) of miRNA profiles of hESC, DE, Duo and Ile spheroids. (D) Heatmap showing changing expression of miRNAs across hESC, DE, Duo and Ile spheroids. Only showing those miRNAs with RPMMM > 1000 in at least one stage. (E-G) Volcano plots showing differentially expressed miRNAs in the event of DE formation (F), SI lineage specification (F), and SI regional patterning (G). Numbers in bronze and pine are for up- and down-regulated miRNAs in the indicated comparison (RPMMM > 1000 in at least in one stage, log2FC > 0.5, padj < 0.05 by Wald test; DESeq2). RPMMM, reads per million mapped to miRNAs. smRNA-seq: human embryonic stem cell (hESC), n = 3; definitive endoderm (DE), n = 3; Duo spheroid (Duo), n = 6; Ile spheroid (Ile), n = 4.

By performing differential expression analysis (see miRNA lists in **Supplementary Data 3**), we identified 35 miRNAs altered during DE formation (16 down and 19 up; **Figure 1E**), 61 miRNAs altered during SI lineage specification (26 down and 35 up; **Figure 1F**), and 49 miRNAs altered during distal SI patterning (23 down and 26 up; **Figure 1G**). The dramatically changing miRNA landscape during directed differentiation towards SI lineage is suggestive of active miRNA-mediated gene regulation during early SI development.

### Integration of ChRO-seq and RNA-seq defines post-transcriptionally regulated genes during major stages of early human SI development

In theory, the depletion and enrichment of miRNAs during a stage transition should attenuate and enhance, respectively, post-transcriptional suppression of target genes. Therefore, we sought to identify genes that are subject to significant post-transcriptional (PT) regulation during each stage transition of the hPSC-HIO model by generating and integrating ChRO-seq and RNA-seq data. Given that ChRO-seq measures nascent transcriptional activity and RNA-seq measures steady-state expression levels of genes, the difference between the two sequencing readouts represents PT regulation (**Figure 2A**). We performed two-factor DESeq2 analysis to identify genes that exhibit discordant changes at the level of transcription (ChRO-seq) versus steady-state expression (RNA-seq) in a given stage transition (**Figure 2B**; **Methods**). For each stage transition, we identified genes for which PT suppression is lost (denoted hereafter as “stable genes”) or genes for which PT suppression is gained (denoted hereafter as “unstable genes”) (two factor analysis: log2FC > 0.5 or < −0.5, padj < 0.2 in Wald test DESeq2; **Figure 2C-E**, see gene lists in **Supplementary Data 4**). This analysis revealed a total of 825 stable and 612 unstable genes during DE formation from hESC (**Figure 2C**); 1972 stable and 1756 unstable genes during SI lineage specification from DE to Duo (**Figure 2D**); and 505 stable and 609 unstable genes during distal SI patterning from Duo to Ile (**Figure 2E**). We further pruned these gene lists by applying a specific stringent criterion: very little change in nascent transcriptional activity (padj > 0.1 in ChRO-seq) but significant alterations in steady-state expression levels (log2FC > 0.5 or < −0.5, padj < 0.05 in RNA-seq). This stringent set signifies genes (hereafter as “stringent stable” or “stringent unstable”) that are regulated primarily through post-transcriptional mechanisms during one or more stage transitions (**Figure 2B**, see gene lists in **Supplementary Data 4**). This analysis revealed 236 stringent stable and 208 stringent unstable genes during DE formation (**Figure 2F**); 292 stringent stable and 324 stringent unstable genes during SI lineage specification (**Figure 2G**); and 139 stringent stable and 139 stringent unstable genes during distal SI patterning (**Figure 2H**). For the rest of the study, we focused mostly on the initial event of SI lineage specification: the transition from DE to Duo, during which we observed the most remarkable changes in the miRNA landscape (**Figure 1B**) as well as the greatest number of stable/unstable genes (which is indicative of the largest amount of PT regulation) (**Figure 2D**).

**Figure 2.**
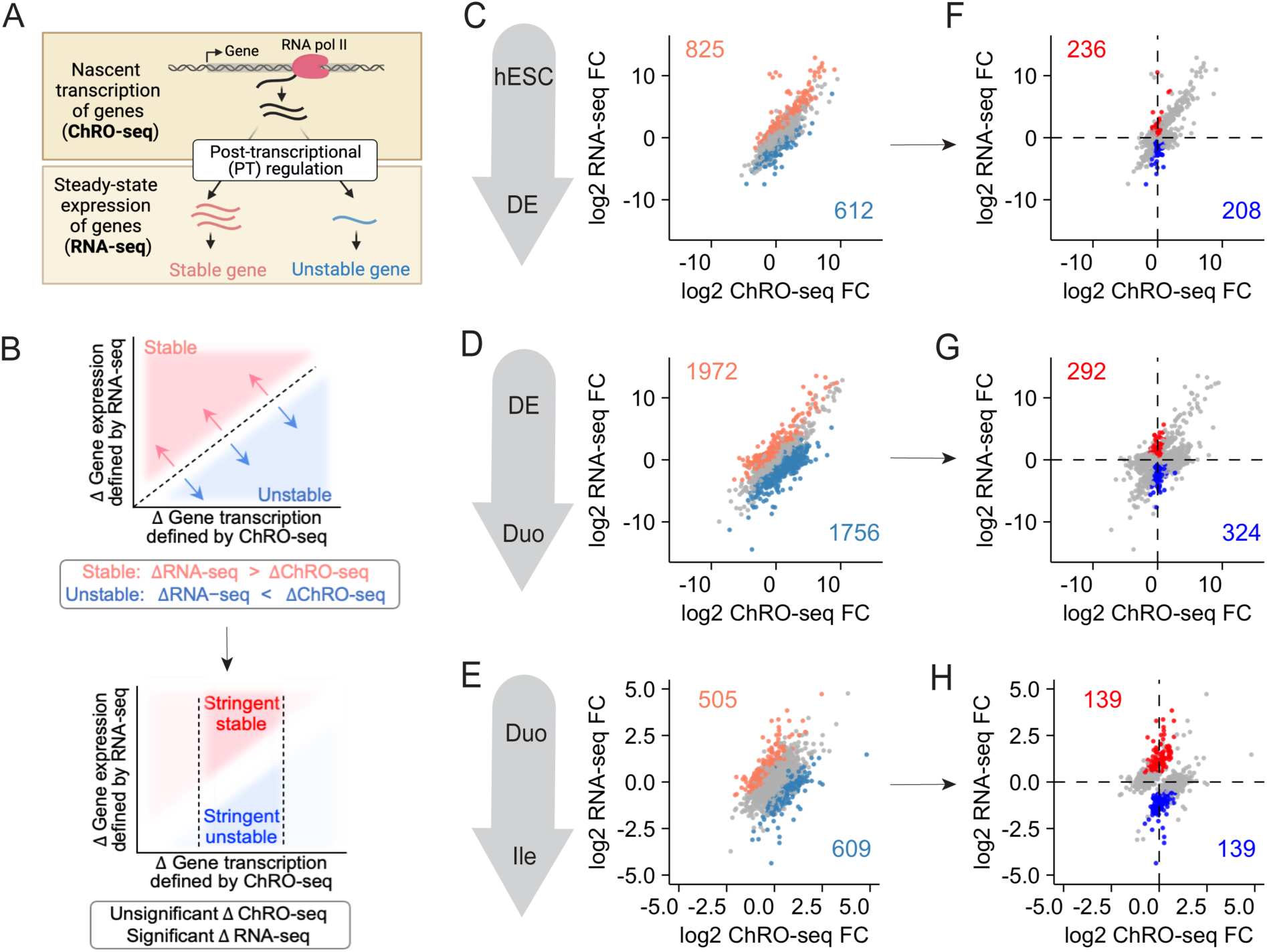
Integrative analysis of ChRO-seq and RNA-seq defined genes that were post-transcriptionally (PT) regulated during early SI development in a human organoid model. (A) Integration of ChRO-seq and RNA-seq defined genes that were PT stable and unstable during a stage transition. (B) Top: Discordant RNA-seq and ChRO-seq changes were detected by two-factor DESeq2 analysis (padj < 0.05 and log2FC > 0.5 in two-factor DESeq2; ChRO-seq TPM > 20; RNA-seq normalized counts > 100). Bottom: Additional filtering (padj > 0.1 in DESeq2 of ChRO-seq; padj < 0.05, log2FC > 0.5 in DESeq2 of RNA-seq) was applied to define genes that were primarily subject to PT regulation during a stage transition. (C-E) Two-factor DESeq2 analysis defined PT regulated genes during the event of DE formation (C), SI lineage specification (D) and SI regional patterning (E). In (C-E), genes in coral and teal are defined as stable and unstable genes, respectively. (F-H) Additional filtering defined genes that were primarily subject to PT regulation during the event of DE formation (F), SI lineage specification (G) and SI regional patterning (H). In (F-H), genes in blue and red are defined as stringent stable and unstable genes, respectively. ChRO-seq: human embryonic stem cell (hESC), n = 3; definitive endoderm (DE), n = 4; Duo spheroid (Duo), n = 3; Ile spheroid (Ile), n = 3. RNA-seq: hESC, n = 2; DE, n = 3; Duo, n = 6; Ile, n = 4.

### Elevated activity of miR-182 leads to post-transcriptional suppression of *SOX2* during SI lineage specification

A miRNA can serve as a master regulator of gene expression in various contexts by targeting key regulators atop the regulatory hierarchy, such as transcription factors (TFs). The stage transition from DE to Duo spheroids is driven by the induction of CDX2, which is required for SI lineage specification and is therefore known as the master TF of intestinal identity (34, 35). Concomitant with acquisition of SI lineage specification is the suppression of DE associated TFs (e.g., SOX17, GATA6, EOMES, LHX1)(36-40), and the suppression of SOX2, which is essential for foregut identity (41, 42). During SI lineage specification (DE to Duo), we observed concordant RNA-seq and ChRO-seq changes in *CDX2, SOX17* and *GATA6*, indicating that the steady-state expression changes in these genes are driven primarily by the changes in nascent transcriptional activity (**Figure 3A**). In contrast, the ChRO-seq and RNA-seq changes are discordant for *EOMES, LHX1* and *SOX2*; specifically, there are no remarkable alterations in nascent transcription at these gene loci, and yet the steady-state mRNA levels are significantly reduced. This finding is indicative of extensive post-transcriptional suppression (**Figure 3A**). Indeed, we defined *EOMES* and *LHX1* as unstable genes (**Figure 2C**) and *SOX2* as a stringent unstable gene (**Figure 2F**) during SI lineage specification.

**Figure 3.**
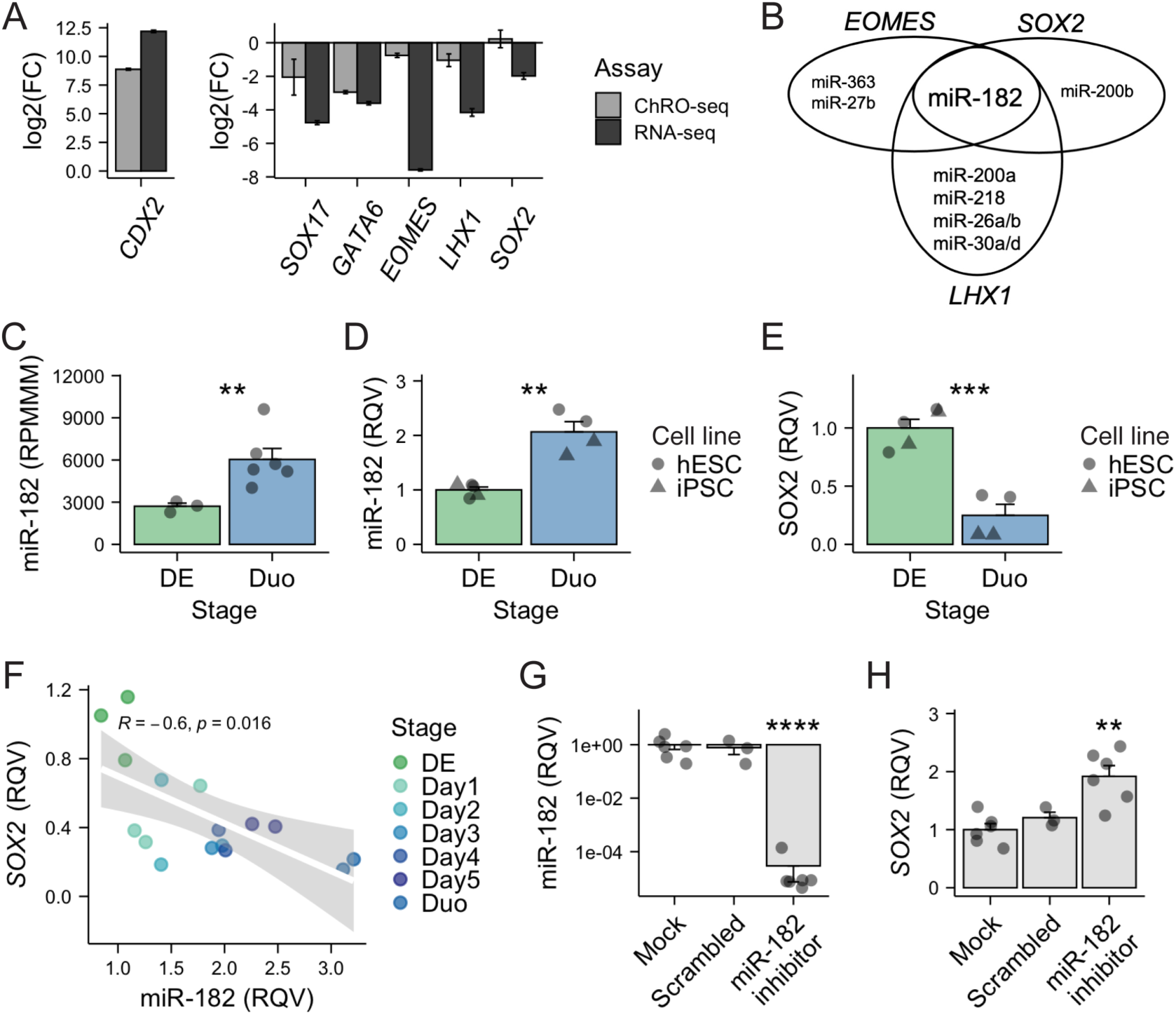
Elevated activity of miR-182 promotes SI lineage formation in part through post-transcriptionally suppressing *SOX2*. (A) Nascent transcriptional (ChRO-seq) and steady-state expressional (RNA-seq) changes of transcription factors critical for SI lineage specification (transition from DE to Duo). (B) miRNAs that are enriched during SI lineage specification and have conserved target sites on *EOMES, LHX1* and *SOX2*. (C) smRNA-seq showing a significant upregulation of miR-182 during SI lineage specification. DE, n = 3; Duo, n = 6. (D) qPCR showing a significant upregulation of miR-182 during SI lineage specification in both hESC and iPSC cell line. N = 2-3 samples per cell line per stage. (E) qPCR showing a significant downregulation of *SOX2* during SI lineage specification in both hESC and iPSC cell line. N = 2-3 samples per cell line per stage. (F) Reverse Spearman correlation between miR-182 and *SOX2* during 5-day window of DE cells receiving SI induction cocktail. N = 2-3 samples per day. (G) qPCR of miR-182 levels in iPSC directed differentiated cells receiving treatment of mock, scrambled control or miR-182 inhibitors. (H) qPCR of *SOX2* (J) in iPSC directed differentiated cells receiving treatment of mock, scrambled control or miR-182 inhibitors. N = 3-6 per condition across 2 independent experiments. DE, definitive endoderm. Duo, duodenal spheroid. RPMMM, reads per million mapped to miRNAs. RQV, relative quantitative value.

To explore candidate miRNAs that mediate the post-transcriptional suppression of *EOMES, LHX1* and *SOX2*, we identified miRNAs that are elevated during this stage transition (**Figure 1E**) and also have highly conserved target sites in these genes. This analysis revealed only one miRNA: miR-182 (**Figure 3B**). In addition, among these three TFs, SOX2 is the only one that satisfies the criteria for “stringent unstable gene” (**Figure 2G**). We next sought to support miR-182-mediated suppression of *SOX2* during SI lineage specification. First, we validated the up-regulation of miR-182 and suppression of *SOX2* during the transition from DE to Duo spheroid in two independent experiments using two different hPSC lines (**Figure 3C-E**). Second, we demonstrated a strong negative correlation between miR-182 and *SOX2* over the course of the 5-day SI fate specification process (Pearson R = −0.6, p = 0.016; **Figure 3F**).

Next, to test miR-182 regulation of *SOX2* during SI lineage specification, we performed miR-182 loss-of-function studies during the transition from DE to Duo spheroids. Specifically, transfection of miR-182 inhibitors (500 nM) was performed in DE cells on day 3 of SI induction and the cells were collected on day 4, the earliest time point at which Duo spheroids are formed in the model. As expected, the inhibition of miR-182 led to a robust suppression of miR-182 compared to the mock and scrambled control groups (**Figure 3G**). While 24-hour treatment with miR-182 inhibitors was not sufficient to reduce CDX2 and block SI fate (**Supplementary Figure 2**), we did observe significant elevation of *SOX2* compared to mock and scrambled control groups (**Figure 3H**. These findings indicate that the up-regulation of miR-182 is one component of a larger regulatory network that promotes SI lineage induction in part through targeting *SOX2*.

### Stable genes during SI lineage specification are strongly associated with the depletion of miR-375

To determine which if any miRNAs are likely most responsible for post-transcriptional regulation during SI lineage specification, we leveraged our previously developed tool called miRhub (43) that determines enrichment of miRNA target sites in a given gene set. We found that stringent unstable genes (defined in **Figure 2G**) during the stage transition from DE to Duo are significantly enriched for predicted target sites of six miRNAs that are significantly up-regulated during the same transition (miR-24, miR-34a, miR-423, miR-26a, miR-30d, miR-744) (**Supplementary Figure 3B**). We also found that stringent stable genes (**Figure 2F**) during the same stage transition are significantly enriched for predicted target sites of only two miRNAs that exhibit reduced expression (miR-375 and miR-302a) (**Figure 4A**). Similar analyses were also performed for the other stage transitions in the hPSC-HIO model (**Supplementary Figure 3**).

**Figure 4.**
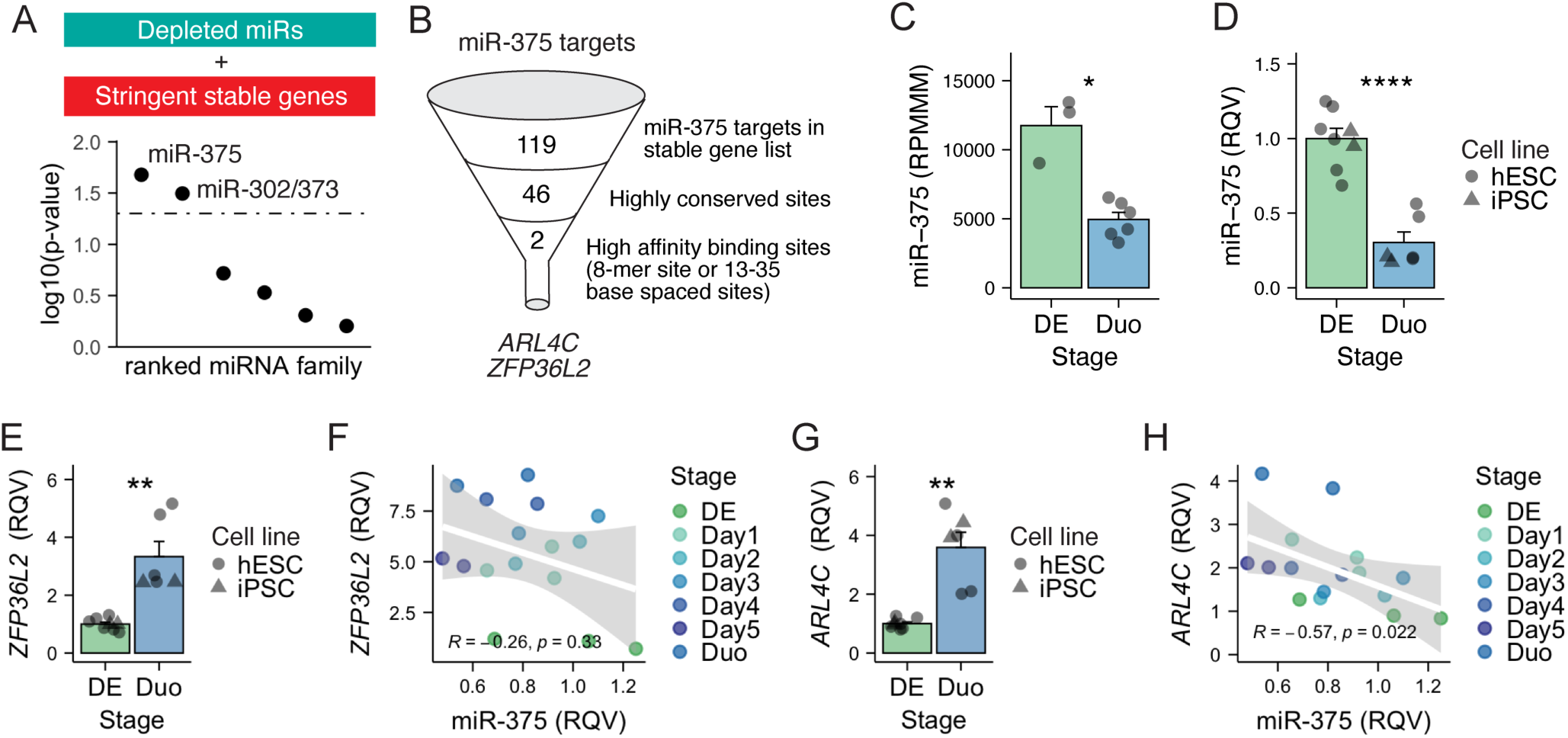
Post-transcriptionally stable genes during SI lineage specification are strongly associated with the depletion of miR-375. (A) MiRhub analysis suggested that genes that stabilized during SI lineage specification were enriched in target sites for two depleted miRNAs, miR-375 and miR-302. (B) miR-375 target genes (n = 119; TargetScan) among stable genes in SI lineage specification. (C) smRNA-seq showing a significant reduction of miR-375 during SI lineage specification. DE, n = 3; Duo, n = 6. (D) qPCR showing a significant reduction of miR-375 during SI lineage specification in both hESC and iPSC cell line. N = 2-4 samples per cell line per stage. (E) qPCR showing a significant downregulation of *ZFP36L2* during SI lineage specification in both hESC and iPSC cell line. N = 2-4 samples per cell line per stage. (F) Modest reverse Spearman correlation between miR-375 and *ZFP36L2* during 5-day window of DE cells receiving SI induction cocktail. N = 2-3 samples per day. (G) qPCR showing a significant downregulation of *ARL4C* during SI lineage specification in both hESC and iPSC cell line. N = 2-4 samples per cell line per stage. (H) Significant reverse Spearman correlation between miR-375 and *ARL4C* during 5-day window of DE cells receiving SI induction cocktail. N = 2-3 samples per day. DE, definitive endoderm. Duo, duodenal spheroid. RPMMM, reads per million mapped to miRNAs. RQV, relative quantitative value.

Among the stable genes during the transition from DE to Duo, 119 have validated or predicted target sites for miR-375 (**Figure 4B**, see gene list in **Supplementary Data 5**). Less than half of these (n=46) have at least one miR-375 target site that is conserved across multiple species (**Figure 4B**). Two of these genes namely *ARL4C* and *ZFP36L2*, have particularly high affinity predicted binding sites for miR-375 (**Figure 4D**). Specifically, *ARL4C* harbors an 8-mer site (Watson–Crick match to miRNA positions 2–8 followed by ‘A’), which is thought to be a high affinity match for miR-375 (based on TargetScan prediction) (**Figure 4E**). *ZFP36L2* has two highly conserved, closely spaced (33 bases apart) miR-375 target sites (**Figure 4B**), a scenario that typically leads to a synergistic suppression effect (44, 45). We first demonstrated that the reduction of miR-375 as well as the up-regulation of *ZFP36L2* and *ARL4C* during the transition from DE to Duo are reproducible across independent directed differentiation experiments with two different hPSC lines (**Figure 4C, 4D, 4F**). Our data showed that miR-375 exhibits a modest negative correlation with *ZFP36L2* (R = −0.26, p = 0.33; **Figure 4E)** and a strong negative correlation with *ARL4C* (R = −0.57, p = 0.022; **Figure 4G**).

We next sought to interrogate miR-375-mediated regulation of *ZFP36L2* and *ARL4C*. It is important to note that *ZFP36L2* and *ARL4C* harbor conserved target sites not only for miR-375 but also for miR-302, both of which are depleted during SI lineage specification and identified as strong candidate miRNA regulators of genes that lose PT suppression during SI lineage specification (**Figure 4A**). To assess the regulatory impact of miR-375 apart from miR-302, we turned to adult mouse SI tissues in which miR-302 is absent but miR-375 remains robustly expressed (46).

### Single cell transcriptomics reveals a post-developmental role for miR-375 in regulating *Zfp36l2* and *Arl4c* in adult mouse SI intestinal stem cells

First we generated mice with whole body knockout of miR-375 using CRISPR/Cas9 technology (375KO mice). We then performed single cell RNA-seq of crypts isolated from 375KO and wild-type mice. This experiment captured a total of 5798 cells from SI epithelial tissues (n = 2591 cells from wild-type; n = 3207 cells from 375KO) after quality control filtering (**Figure 5A-C; Supplementary Figure 4**). All major intestinal epithelial cell types were recovered from both wild-type and 375KO tissues (**Supplementary Figure 4**). We observed that *Zfp36l2* is robustly expressed across different cell types present in adult SI epithelium and especially enriched in progenitor and stem cell populations (**Figure 5C-D**). We performed differential expression analysis in 375KO vs. wild-type for each cell type, which demonstrated that the absence of miR-375 leads to the significantly upregulation of *Zfp36l2* in intestinal stem cells (**Figure 5E-F**). Although *Arl4c* is very lowly detected in our single cell RNA-seq data, we performed pseudo-bulk differential expression analysis and found that *Arl4c* is elevated in 375KO relative to wild-type (1.42 fold increase, P = 0.37 by two-tailed t-test; **Supplementary Figure 5**). Collectively, this study provides *in vivo* evidence to support a life-long role for miR-375 in regulating *Zfp36l2*.

**Figure 5.**
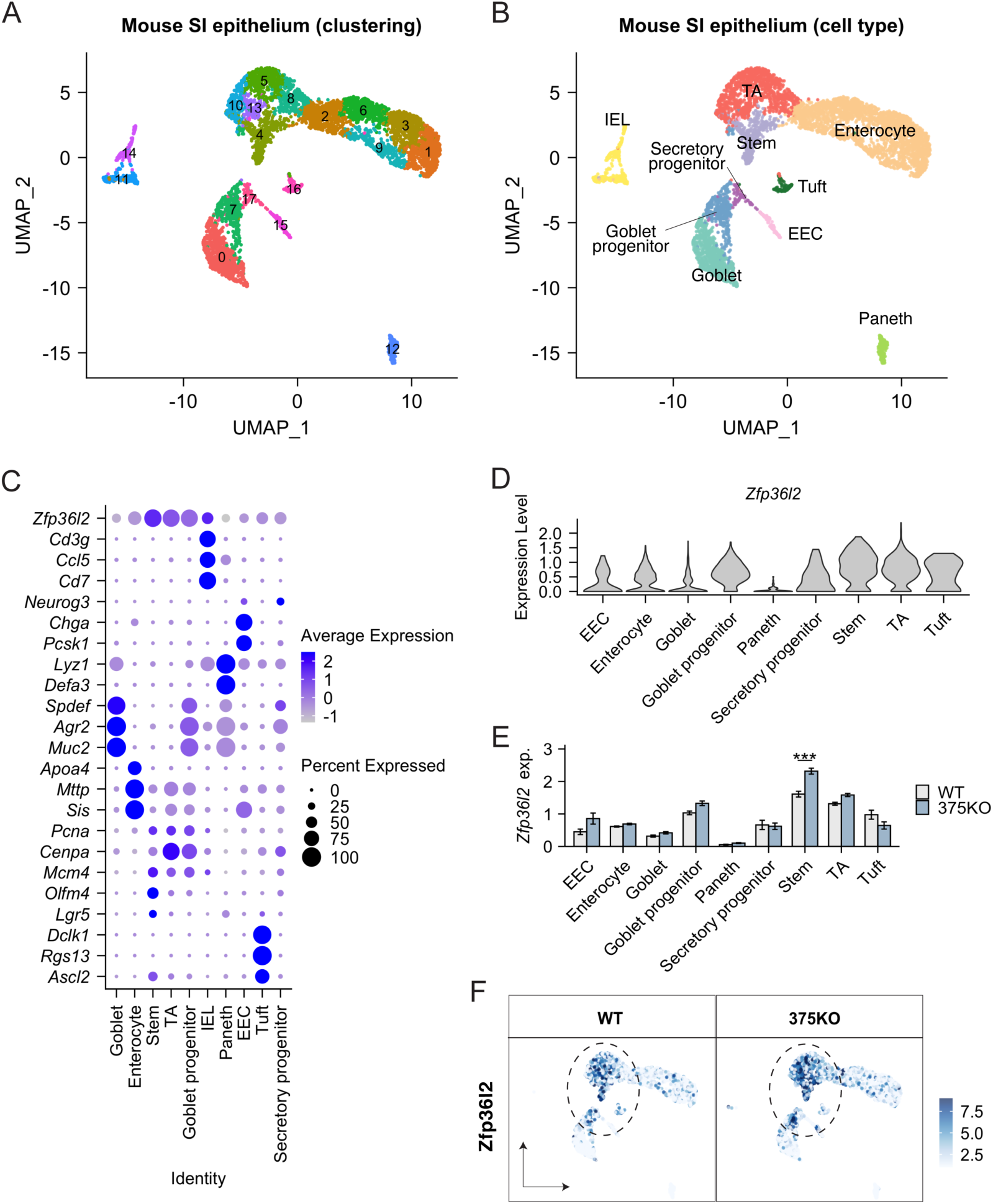
Single cell RNA-seq revealed a long-lasting, conserved role for miR-375 in regulating Zfp36l2 in intestinal stem cell population of adult mice. (A) UMAP of initial clustering analysis with intestinal epithelial cells of adult WT and whole body miR-375 knockout (375KO) mice. N = 2 mice per genotype; 2937 cells from WT and 3275 cell from 375KO after doublet removal and quality control filtering. (B) UMAP of cell type annotation with intestinal epithelial cells of adult WT and 375KO mice. (C) Dot plot of known cell type markers and *Zfp36l2* across all cell types defined in the single cell RNA-seq dataset. (D) Violin plot showing *Zfp36l2* most enriched in stem cell population. (E) Significant upregulation of *Zip36l2* in stem cell population of 375KO compared to WT mice (adjusted P value = 0.000279; log2FC = 0.346). (F) UMAP overlain with the normal expression of *Zfp36l2* in WT and 375KO mice. UMAP, Uniform Manifold Approximation and Projection. IEL, Intraepithelial lymphocytes; EEC, enteroendocrine cell; TA, transient amplifying cell.

### SMAD4 and WNT signaling exert opposite effects on regulating miR-375 expression

Because our data suggests that up-regulation of miR-182 and reduction of miR-375 are critical events during SI lineage specification (**Figure 3-4**), we were motivated to understand how the expression of these miRNAs are regulated. We found that the upregulation of miR-182 during SI lineage specification is likely driven by stabilization of miR-182 transcripts, as transcriptional activity of the miR-182 locus was unchanged whereas smRNA-seq showed a robust increase in mature miR-182 levels during the DE-to-Duo transition (**Figure 6A**). On the other hand, we found that the depletion of mature miR-375 levels (smRNA-seq) is matched by the reduction in nascent transcription (ChRO-seq), indicating that miR-375 is regulated during the DE-to-Duo transition largely at the transcriptional level (**Figure 6A**). Indeed, we observed high transcriptional activity around the miR-375 locus only in the DE stage, but not in other stages including Duo spheroids (**Figure 6B**). We mined previously defined transcriptional regulatory elements (TREs) in the hPSC-HIO model (26) and found that four out of the six TREs adjacent to the miR-375 locus (TRE#2, #3, #4 and #6) are uniquely active in DE (**Figure 6B**). We also observed that the summed transcriptional activity of putative enhancer TREs (TRE#1-4) is highly correlated with the transcriptional activity of TREs overlapping with the putative promoter region of miR-375 (TRE#5-6) across all four stages of the directed differentiation model (**Figure 6C**). Taken together, these observations demonstrate that miR-375 transcription is in large part regulated by the activity of nearby enhancers and that the significant down-regulation of mature miR-375 during SI lineage specification is likely due to reduced transcriptional activity at the corresponding genomic locus.

**Figure 6.**
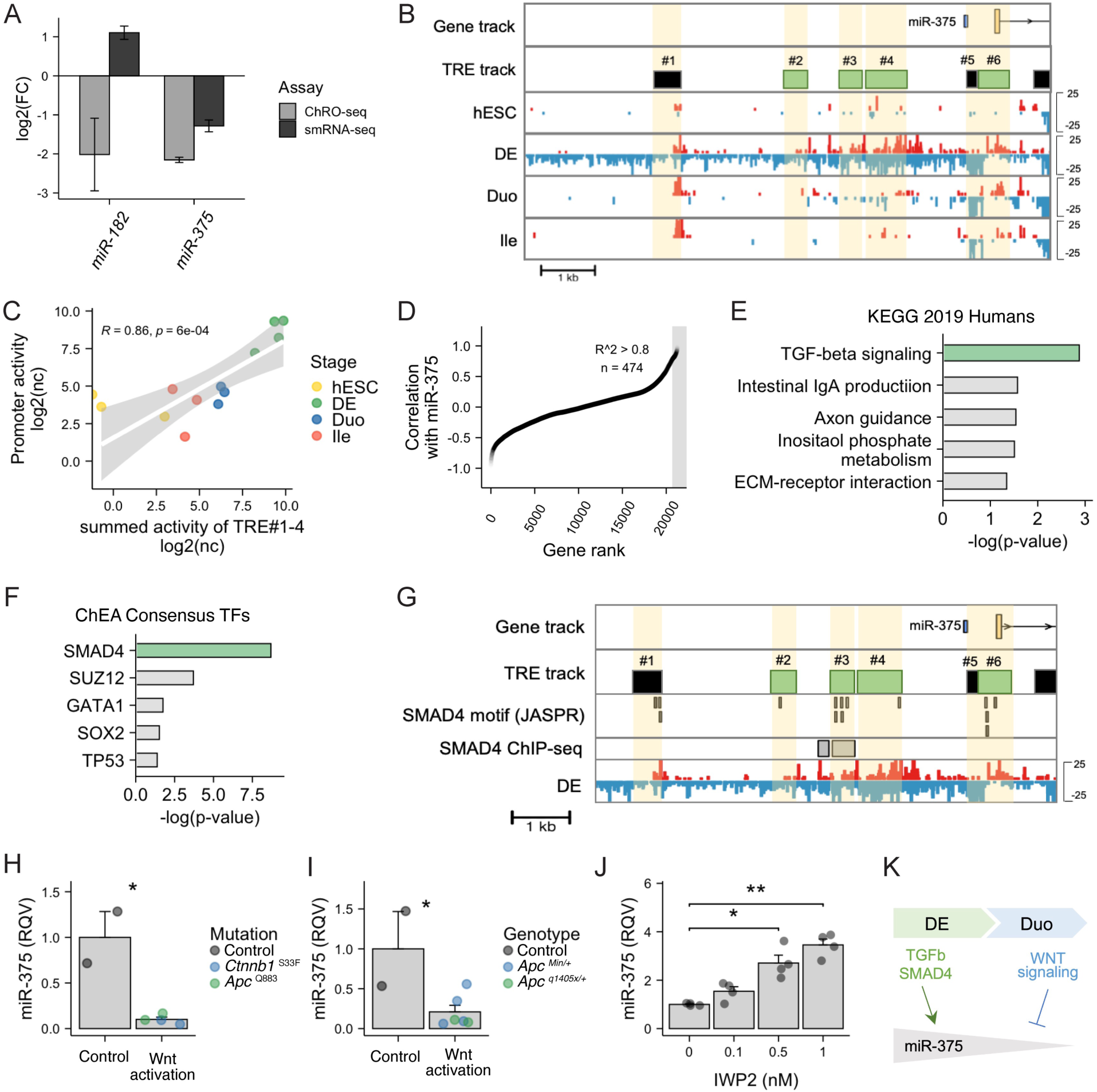
SMAD4 and WNT signaling respectively serve as a positive and negative regulator of miR-375 expression in SI epithelial cells. (A) Nascent transcriptional (ChRO-seq) and steady-state expressional (smRNA-seq) changes of miR-182 and miR-375 during SI lineage specification (transition from DE to Duo). (B) Normalized ChRO-seq signal and active TREs detected around miR-375 locus across different stages. Reads in red and blue are signals from positive and negative strand, respectively. DE-specific TREs (in green) were defined previously in Hung et all., (2021). Scale (± 25 reads/kb/10^6^) is fixed across all stages. (B) Positive Spearman correlation between ChRO-seq activity of putative miR-375 promoter (TRE#5-6) and the summed activity of nearby enhancers (TRE#1-4) across all stages. (D) Genes ranked based on their correlation with miR-375 expression across stages of hESC, DE, Duo spheroid and Duo HIO. Genes with positive correlation with miR-375 expression (n = 474; R > 0.8) are highlighted. (E) Enrichment of TGF beta signaling (KEGG 2019 Human) in genes that have positive correlation with miR-375 (R > 0.8). (F) Enrichment of SMAD4 (ENCODE & ChEA consensus) in genes that have positive correlation with miR-375 (R > 0.8). (G) DE-specific TREs that are nearby miR-375 locus contain sequences of SMAD4 motif (MA1153.1; JASPAR) and overlap with SMAD4 sites defined by the ChIP-seq study (Kim et al., 2011). (H) qPCR showing *ex vivo* downregulation of miR-375 in mouse enteroids with mutation-induced Wnt activation (n = 4 including 2 samples with *Apc*^Q883^ and 2 samples with *Ctnnb1*^S33F^ mutation) compared to wild type control (n = 2). *P < 0.05 by two-tailed t-test. (J) qPCR showing *in vivo* downregulation of miR-375 in polyps of *Apc* mutant mice (n = 6 including 4 samples from *Apc*^min/+^ mice and 2 samples from *Apc*^q1405x/+^ mice) compared to the villi scrappings of control group (n = 2). *P < 0.05 by two-tailed t-test. (K) qPCR showing an upregulation of miR-375 in mouse enteroids treated with IWP2, a Wnt signaling inhibitor, compared to mock. N = 4 wells per condition. *P < 0.05, **P < 0.01 by two-tailed t-test. (L) Proposed model of TGFb/SMAD4 and Wnt signaling in regulating miR-375 expression. TRE, transcriptional regulatory element. hESC, human embryonic stem cell. DE, definitive endoderm. Duo, duodenal spheroid. Ile, ileal spheroid. APC, *Adenomatous polyposis coli*.

To explore the candidate pathway(s)/regulator(s) that control miR-375 transcription during human SI development, we first identified 474 genes (RNA-seq data) that are positively correlated (R^2^ > 0.8) with miR-375 expression across hESC, DE, Duo spheroid (Day 0 HIOs) and Duo HIO (Day 28 HIOs) (**Figure 6D**). Pathway analysis of these genes revealed significant enrichment of TGFb signaling (P = 0.0013 in KEGG 2019 human database; **Figure 6E**) and transcriptional networks controlled by SMAD4 (P = 2.046544e-09 in ChEA Consensus TFs database; **Figure 6F**). Consistent with these findings, we observed that DE-specific TRE#3 harbors several high-confidence SMAD4 motifs (JASPAR database), which have been validated by SMAD4 ChIP-seq in hESC-derived DE (47)(**Figure 6G**). These findings together suggest key roles for SMAD4 in promoting transcription at the miR-375 locus (**Figure 6L**).

During SI lineage specification, the levels of miR-375 are significantly reduced upon treatment with a cocktail that augments Wnt signaling. This motivated us to test whether Wnt signaling itself serves as a negative regulator of miR-375 expression in the SI. To this end, we leveraged CRISPR-engineered mouse enteroids in which a mutation was introduced to *Apc* (*Apc*^*Q883*^) or *Ctnnb1* (*Ctnnb1*^S33F^), leading to constitutive activation of Wnt signaling (48, 49). By performing qPCR analysis, we showed a very robust reduction of miR-375 expression in the Wnt-activated mouse enteroids compared to the control group (∼10 fold downregulation; **Figure 6H**). Consistent with this finding, we observed a significant reduction of miR-375 in SI polyps developed in mouse models with hyperactivation of Wnt signaling (*Apc*^Min/+^ or *Apc*^q1405x/+^)(48), compared to control mice (**Figure 6I**). We also showed that treatment of WT mouse enteroids with Wnt signaling inhibitor, IWP2, significantly elevated miR-375 expression in a dose-dependent manner (**Figure 6J**). Taken together, these data strongly support Wnt signaling as a negative regulator of miR-375 expression during SI lineage specification (**Figure 6K**).

## Discussion

The prenatal development of the SI requires a well-orchestrated transcriptional program to achieve cellular identity acquisition and proper morphological patterning (50, 51). The present study represents the first investigation to our knowledge of miRNA-mediated post-transcriptional mechanisms in very early events of human SI development. An important SI developmental event included in this study is SI lineage specification from the endodermal layer, which occurs during post-conceptual week 2 of human development. Studying such early developmental events in humans remains extremely challenging due to limited access to primary human fetal tissues from appropriate embryonic stages. A major strength of this study is the use of the directed differentiation model that generates fetal-like human intestinal organoids (HIOs) from human pluripotent stem cells (hPSCs). By leveraging this state-of-art human model of SI development, we provide novel insights into post-transcriptional regulation and miRNA activity during human SI development.

Our smRNA-seq analysis demonstrated changing miRNA signatures across different cellular stage transitions of the hPSC-HIO model. The up- or down-regulation of miRNAs during a cellular stage transition can in theory lead to changes in expression of target genes. Leveraging RNA-seq data alone to test this hypothesis is fraught with limitations, as changes detected in steady-state gene expression levels by RNA-seq could be due to transcriptional and/or post-transcriptional (PT) processes. ChRO-seq was recently developed not only to detect active regulatory elements but also provide sensitive measurements of gene transcriptional activity (25, 26). In this study, by integrating RNA-seq and ChRO-seq data, we teased apart transcriptional from PT regulation of genes across the entire genome and identified potential miRNA regulators relevant to human SI development.

We showed how a miRNA may exert gene regulatory impact through targeting an important TF during early SI development. Specifically, we found that the increase in miR-182 leads to the suppression of *SOX2*. Although miR-182 has been shown to target and regulate *SOX2* previously in cochlea development and glioblastoma (52, 53), our study demonstrated the functional relevance of this regulatory connection in the context of SI development. SOX2 is known as a master TF of foregut identity (41, 42); therefore, the increased expression and activity of miR-182 during SI lineage establishment may serve as a critical mechanism to favor midgut vs. foregut lineages.

We also demonstrated that the depletion of miR-375 during SI lineage specification significantly alleviates post-transcriptional suppression and thereby upregulates its target genes, including *ARL4C* and *ZFP36L2*, which could play an important role during this developmental event. ARL4C, which belongs to the superfamily of GTP-binding proteins, is reported to control the formation of 3D tubular structure in the rat intestinal IEC6 cell line (54). *ZFP36L2* is a previously validated miR-375 target in pancreatic carcinoma (55) and known to code for an RNA binding protein that can destabilize RNA transcripts containing AU-rich elements (56). Notably, *SOX2* is identified as one of the PT unstable genes during SI lineage formation in our analyses and known to harbor AU-rich elements. Whether *SOX2* is a direct target of ZFP36L2 for mRNA decay during this developmental event warrants further investigation. Both *ZFP36L2* and *ARL4C* also harbor conserved predicted target sties for miR-302, which our analysis suggests may work in coordination with miR-375 to regulate SI lineage specification. To delineate the regulatory impact of miR-375 apart from that of miR-302, we studied adult mouse SI tissue in which miR-302 is absent but miR-375 is robust expressed. As shown by our scRNA-seq study of adult miR-375KO mice, the miR-375:*Zfp36l2* and miR-375:*Arl4c* regulatory interactions are conserved across species and maintained in adult intestinal stem cells beyond early SI development. This long-lasting regulatory impact of miR-375 could in part contribute to changes in intestinal function and tissue homeostasis observed previously (57, 58).

Our study also revealed that while miR-182 expression may be regulated primarily by post-transcriptional mechanisms, miR-375 levels are changed largely by nascent transcriptional activity. Our analyses suggested SMAD4 and TGFb signaling as positive regulators of miR-375 expression during human SI development, supporting only one previous study in which it was shown that TGFb/Smad pathway promotes miR-375 expression in pancreatic islets (59). By mining our ChRO-seq data, we showed that transcription of miR-375 is likely driven by nearby enhancers, one of which harbors a previously validated SMAD4 binding site (47). In addition, our data suggests that miR-375 and WNT signaling have a reciprocal inhibitory relationship. Many of the miR-375 targets identified in our analyses are potentiators of WNT signaling (e.g., *ARL4C, ZFP36L2, TCF12, TCF4*). We also demonstrated that WNT signaling is a robust negative regulator of miR-375.

Many studies have uncovered individual miRNAs that control certain aspects of adult SI biology, including SI cell type allocation and function (60-63), epithelial homeostasis (57, 64-66), and disease pathogenesis (67-69). The present study uncovers a novel facet of miRNAs in regulating prenatal SI development. We leveraged multi-omic, systems biology approaches to discover candidate miRNA regulators associated with the events of DE formation, SI lineage specification, and SI regional patterning in a human organoid model. While this work mainly focused on post-transcriptional regulation relevant to SI lineage specification, the candidate miRNA regulators that we identified for the other stages of SI development also warrant detailed characterization in the future.

## Methods

### Directed differentiation of human pluripotent stem cells

The human embryonic stem cell (hESC) line H9 (NIH registry #0062) was obtained from the WiCell Research Institute. The induced pluripotent stem cell line (iPSC) 72.3 was obtained from Cincinnati Children’s Hospital (70). hPSCs were maintained and passaged in mTeSR1 medium (STEMCELL Technologies, Vancouver, BC, Canada) in cell culture plates coated with Matrigel (BD Biosciences, San Jose, CA). hPSCs were differentiated toward human duodenal or ileal fates as previously described (22, 23, 27). Briefly, the definitive endoderm was generated by treatment of activin A (100 ng/ml) for 3 consecutive days in Roswell Park Memorial Institute 1640 (RPMI-1640) media supplemented with 0% (v/v; day 1), 0.2% (v/v; day 2) and 2.0% (v/v; day 3) Hyclone defined fetal bovine serum (dFBS). Midgut patterning that generates intestinal spheroids was carried out in RPMI-1640 supplemented with 2% HyClone dFBS, FGF4 (500 ng/mL) and CHIR99021 (2 μM) with daily media changes. The intestinal spheroids that represent progenitor cells of human fetal duodenum and ileum were collected after 5 and 10 days of FGF4/CHIR99021 cocktail, respectively.

### Whole body miR-375 knockout mouse model

Mouse miR-375 gene was deleted using the CRISPR/Cas9 system with a guide RNA (gRNA) targeting the 946-965 bp region of the mouse miR-375 gene (ENSMUSG00000065616). 173 FVB x B62J F1 hybrid 1-cell embryos were co-injected with 2.5 μg of gRNA and 7.5 μg of Cas9 mRNA. 107 embryos at two-cell stage were transferred to pseudo-pregnant recipient female mice at ∼20 embryos/recipient. 20 founder pups were born and validated for deletion of the miR-375 gene by targeted genomic sequencing. Mice null for miR-375 were backcrossed at least three generations with B6(Cg)-Tyrc-2J (B6 albino) wildtype mice. Synthesis of the gRNA, mouse embryo co-injections and viable embryo emplacement were performed by the Cornell Stem Cell and Transgenic Core Facility at Cornell University and supported in part by Empire State Stem Cell Fund (contract number C024174). Genotyping was performed on genomic DNA extracted from tail tissues, using forward primer 5′-GAAGCTCATCCACCAGACACC-3′ and reverse primer 5′-GAGCTATGCCTGGCAGCTCA-3′ amplifying a 307 bp product in wild-type mice and a 260 bp product in miR-375 knockout mice. All animal procedures were performed with the approval and authorization of the Institutional Animal Care and Use Committee at Cornell University.

### *Apc* mutant mouse models

Generation and genotype validation of *Apc*^Min/+^ and *Apc*^q1405x/+^ mouse line were Described previously in (48). These mouse lines with a heterozygous mutation at *Apc* locus have been shown to form polyps in the small intestine due to a hyperactivation of Wnt signaling (48). In this study, small intestinal polyps from *Apc*^Min/+^ and *Apc*^q1405x/+^ mice were identified using a dissection microscope, individually collected from cold PBS-flushed small intestine, and subsequently flash frozen at −80°C. Collected villi scrappings of Wild type mice from Rosa26-CAGs-LSL-rtTA3 knock-in or Rosa26-CAGs-rtTA3 knock-in colonies were used as control group. All animal procedures were performed with the approval and authorization of the Institutional Animal Care and Use Committee at Weill Cornell Medicine.

### Isolation of mouse jejunal crypts

Jejunal crypts were isolated from male C57BL/6 or B6 albino mice as previously described (71). Small intestine was excised from mice and divided into three equal segments. The middle region was considered jejunum. Subsequent to luminal flushing with ice cold PBS, the tissue was longitudinally cut and subjected to incubation in 3 mM EDTA in ice cold PBS with 1% (v/v) Primocin™ (InvivoGen, San Diego, CA; ant-pm-1) for 15 min at 4°C. The mucosa of the intestinal pieces was gently scrapped of mucus, shaken in ice cold PBS with 1% (v/v) primocin for 2 minutes, and incubated in fresh 3 mM EDTA in ice cold PBS with 1% (v/v) primocin for 40 min at 4°C. After 2 to 6 minutes of gentle manual shaking in ice cold PBS with 1% (v/v) primocin, the intestinal pieces were inspected microscopically for detached intestinal crypts, diluted 1:2 with ice cold PBS with 1% (v/v) primocin, filtered with a 70 μm cell strainer, and collected by pelleting with centrifugation at 110 g for 10 min at 4°C. The isolated crypts were used for enteroid culture or single cell RNA-seq library preparation.

### Mouse enteroid culture

Mouse enteroid culture was performed as previously described (57, 64). The jejunal crypts (day 0) isolated from male 3-to 5-month-old B6 albino mice were grown in Reduced Growth Factor Matrigel (Corning, Corning, NY; 356231) and advanced DMEM/F12 (Gibco; 12634-028) containing GlutaMAX (Gibco; 35050-061), Primocin (Invivogen, San Diego, CA; ant-pm-05), HEPES (Gibco; cat. 15630-080), N2 supplement (Gibco; cat. 17502-048), 50 ng/μL EGF (R&D Systems, Minneapolis, MN; 2028-EG), 100 ng/μL Noggin (PeproTech, Rocky Hill, NJ; 250-38), 250 ng/μL murine R-spondin (R&D Systems, Minneapolis, MN; 3474-RS-050), and 10 μM Y27632 (Enzo Life Sciences, East Farmingdale, NY; ALX270-333). For Wnt suppression studies, IWP2 (Tocris Bioscience, Bristol, UK; 3533), the compound that inhibits Wnt processing and secretion, was add at on day 0 and day 3. Enteroids on day 5 were harvested for total RNA isolation.

### CRISPR/Cas9 editing of mouse enteroids

The proximal 15 cm of the small intestine was harvested from 6-week-old C57BL/6 mice, followed by crypt isolation as previously described (72). Crypts were plated in Reduced Growth Factor Matrigel (Corning, Corning, NY; 356231) and grown in basal media containing 50 ng/μL EGF (Invitrogen Waltham, MA; PMG8043), 100 ng/μL Noggin (PeproTech, Rocky Hill, NJ; 250-38) and 500 ng/μL murine R-spondin (R&D Systems, Minneapolis, MN; 3474-RS-050) for 3-4 weeks (48). To generate enteroids with constitutive Wnt activation, enteroids were dissociated and transfected with using CRISPR base editing to introduce a *Apc* mutation (*Apc*^Q883*^) or *Ctnnb1* mutation (*Ctnnb1*^S33F^) to the system, detailed in (48, 49). Exogenous R-spondin1 were withdrawn 2 days after transfection to select for mutant enteroids.

### Real-time quantitative PCR

Total RNA was isolated using the Total Purification kit (Norgen Biotek, Thorold, ON, Canada) per manufacturer’s instructions. Reverse transcription of miRNA was performed using TaqMan microRNA Reverse Transcription kit (Thermo Fisher Scientific, Waltham MA). miRNA expression was measured using TaqMan assays (Thermo Fisher Scientific, Waltham MA) with TaqMan Universal PCR Master Mix. The microRNA TaqMan assays used in this study are miR-375 (assay ID: 000564) and miR-182 (assay ID: 002334). miRNA expression was normalized to U6 (assay ID: 001973) or RNU48 (assay ID: 001006). Reverse transcription of RNA was performed using High Capacity RNA-to-cDNA kit (Thermo Fisher Scientific, Waltham, MA). Gene expression was measured using TaqMan assays (Thermo Fisher Scientific, Waltham MA) with TaqMan Gene Expression Master Mix. The Gene expression TaqMan assays used in this study are *SOX2* (Hs04234836_s1), *ARL4C* (Hs00255039_s1), *ZFP36L2* (Hs00272828_m1), *CDX2* (Hs01078080_m1) and *SOX17* (Hs00751752_s1) and CDH1 (Hs01023895_m1). RNA expression levels were normalized to RPS9 (Hs02339424_g1). Reaction was quantified on a BioRad CFX96 Touch Real Time PCR Detection System (Bio-Rad Laboratories, Richmond, CA, United States).

### Transfection studies in directed differentiation model

We used human induced pluripotent stem cell line 72.3 for transfection studies. To test candidate miRNA regulators, DE-patterned cells were transfected by miRCURY LNA inhibitors (Qiagen, Hilden, Germany) against specific miRNAs during hindgut fate induction (FGF4/CHIR99021 cocktail). Transfection of LNA inhibitors were performed using Lipofectamine 3000 (ThermoFisher Scientific, Waltham MA; L3000-008) per manufacturer’s instructions. Specifically, on day 3 of FGF4/CHIR99021 cocktail, DE cells were treated with 500 nM LNA miR-182 inhibitor (Qiagen, Hilden, Germany; #4101243-101) or scrambled control (Power Negative Control A; YI00199006-DDA). After 24 hour treatment, cell monolayer was collected for RNA isolation followed by qPCR measurements. The efficacy of directed differentiation was confirmed by qPCR measurements of CDX2 (intestinal marker gene) and SOX17 (DE marker gene) in mock treated cells. The efficacy of LNA transfection was assessed by qPCR of target miRNA.

### Small RNA sequencing

Total RNA was isolated using the Total Purification kit (Norgen Biotek, Thorold, ON, Canada). RNA was quantified with the Nanodrop 2000 (Thermo Fisher Scientific, Waltham, MA), and RNA integrity was determined by the Agilent 2100 Bioanalyzer or 4200 Tapestation (Agilent Technologies, Santa Clara, CA). Libraries were prepared using the CleanTag Small RNA Library Prep kit (TriLink Biotechnologies, San Diego, CA). Sequencing (single-end 50x) was performed on the HiSeq2000 platform (Illumina, San Diego, CA) at the Genome Sequencing Facility of the Greehey Children’s Cancer Research Institute (University of Texas Health Science Center, San Antonio, TX). Mapping statistics of small RNA-seq samples can be found in **Supplementary Data 1**. Raw sequencing and processed data can be accessed through GEO record GSE207467.

### RNA sequencing

Total RNA was isolated using the Total Purification kit (Norgen Biotek, Thorold, ON, Canada). RNA purity was quantified with the Nanodrop 2000 (Thermo Fisher Scientific, Waltham, MA), and RNA integrity was quantified with the Agilent 2100 Bioanalyzer or 4200 Tapestation (Agilent Technologies, Santa Clara, CA). Libraries were generated using the NEBNext Ultra II Directional Library Prep Kit (New England Biolabs, Ipswich, MA, USA) and subjected to sequencing (single-end 92×) on the NextSeq500 platform (Illumina) at the Cornell University Biotechnology Resource Center. At least 80M reads per sample were acquired. Raw sequencing data and miRNA quantification tables can be accessed through GEO record GSE142316.

### Chromatin run-on sequencing

Chromatin isolation for chromatin run-on sequencing (ChRO-seq) was performed as previously described (25, 26). To perform run-on reaction with the isolated chromatin, samples were mixed with an equal volume of 2X run-on reaction mix [10 mM Tris-HCl pH 8.0, 5 mM MgCl2, 1 mM DTT, 300 mM KCl, 400 μM ATP, 0.8 μM CTP, 400 μM GTP, 400 μM UTP, 40 μM Biotin-11-CTP (Perkin Elmer, Waltham, MA; NEL542001EA), 100 ng yeast tRNA (VWR, 80054–306), 0.8 units/μL SUPERase In RNase Inhibitor, 1% (w/v) Sarkosyl]. The run-on reaction was incubated in an Eppendorf Thermomixer at 37°C for 5 min (700 rpm) and stopped by adding Trizol LS (Life Technologies, Carlsbad, CA; 10296–010). RNA samples were precipitated by GlycoBlue (Ambion, Austin, TX; AM9515) and resuspended in diethylpyrocarbonate (DEPC)-treated water. To perform base hydrolysis reaction, RNA samples in DEPC water were heat denatured at 65°C for 40 s and incubated in 0.2N NaOH on ice for 4 min. Base hydrolysis reaction was stopped by neutralizing with Tris-HCl pH 6.8. Nascent RNA was enriched using streptavidin beads (NEB, Ipswich, MA; S1421) followed by RNA extraction using Trizol. RNA samples were then processed through the following steps: (i) 3′ adapter ligation with T4 RNA Ligase 1 (NEB, Ipswich, MA; M0204), (ii) RNA binding with streptavidin beads (NEB, Ipswich, MA; S1421) followed by RNA extraction with Trizol, (iii) 5′ de-capping with RNA 5′ pyrophosphohydrolase (NEB, Ipswich, MA; M0356), (iv) 5′ end phosphorylation using T4 polynucleotide kinase (NEB, Ipswich, MA; M0201), (iv) 5′ adapter ligation with T4 RNA Ligase 1 (NEB, Ipswich, MA; M0204). The 5’ adaptor contained a 6-nucleotide unique molecular identifier (UMI) to allow for bioinformatic detection and elimination of PCR duplicates. Streptavidin bead binding followed by Trizol RNA extraction was performed again before final library construction. To generate ChRO-seq library, cDNA was generated through a reverse-transcription reaction using Superscript III Reverse Transcriptase (Life Technologies, Carlsbad, CA; 18080–044) and amplified using Q5 High-Fidelity DNA Polymerase (NEB, Ipswich, MA; M0491). Finally, ChRO-seq libraries were sequenced (5′ single end; single-end 75×) using the NextSeq500 high-throughput sequencing system (Illumina, San Diego, CA) at the Cornell University Biotechnology Resource Center. Raw and processed sequencing data and can be accessed through GEO upon publication.

### Single cell RNA sequencing

Mouse jejunal crypts from male B6 albino wildtype and miR-375 knockout mice were isolated and resuspended in an ice-cold PBS containing 0.04% (w/v) bovine serum albumin and pelleted at 1000 x g at 4°C for 5 min. The crypts were subsequently digested with 0.3 U/mL Dispase I (MilliporeSigma Burlington, MA; D4818) in HBSS at 37°C for 12 min with gentle agitation. The dispase I activity was stopped by adding fetal bovine serum to a final concentration of 10% (v/v) and DNaseI to a final concentration of 50 μg/mL. The sample was filtered through 40 μm strainer, pelleted at 500 x g at 4°C for 5 min, washed with cold HBSS, filtered again, and then resuspended in an ice-cold PBS containing 0.04% (w/v) bovine serum albumin. Prior to submission, single cell suspensions were triturated by pipetting and evaluated for total viable cell number by using trypan blue staining with a TC20 automated cell counter (Bio-Rad Laboratories, Richmond, CA). Single cell RNA sequencing of these samples was performed at the Cornell University Biotechnology Resource Center. Libraries were prepared using the 10X Genomics Chromium preparation kit (3’ v2 chemistry)(10X Genomics, Pleasanton, CA). Sequencing quality of single cell RNA-seq samples can be found in **Supplementary Data 2**. Raw single cell RNA-seq data of wildtype mice was included in our previous study (GSE178826) and re-analyzed in the present study. Raw sequencing data and process files of miR-375 knockout mice can be accessed through GEO upon publication.

### Small RNA-seq analyses

SmRNA-seq reads were trimmed, mapped, and quantified to the hg19 genome using miRquant 2.0 (73), our previously described smRNA-seq analysis pipeline. Briefly, reads were trimmed using Cutadapt v1.12. miRquant 2.0 requires a minimum 10 nucleotide overlap between the adapter sequence and the 3’-end of the sequencing read with less than 10% errors in the alignment. Those reads that did not meet this criterion, or were < 14 nucleotides in length following trimming, were discarded. Alignment of reads were performed using Bowtie v1.1.0 (74). Perfectly aligned reads represented miRNA loci and imperfectly mapped reads (from miRNA isoforms) were re-aligned to these loci using SHRiMP v2.2.2 (75). If a read aligned equally well to multiple genomic loci, those reads were proportionally assigned to each of the loci. Aligned reads were quantified and normalized using reads per million mapped to miRNAs (RPMMM). Principal components analysis (PCA) of miRNAs was performed using raw counts with rlog transformation. To identify differentially expressed miRNAs, smRNA-seq raw counts were analyzed through DESeq2 v1.30.1(76). MiRNAs that were significantly altered between two conditions were defined by criteria of RPMMM > 1000 at least in one condition, log2 foldchange > 0.5 (or < −0.5) and padj < 0.05 by Wald test (DESeq2).

### RNA-seq analyses

RNA-seq reads were mapped to human genome hg38 using STAR v2.5.3a (77) and transcript quantification was performed using Salmon v0.6.0 (78) with GENCODE v33 transcript annotations. The expression levels of genes were normalized using DESeq2 v1.30.1 (76). To identify differentially expressed genes, RNA-seq raw counts were analyzed through DESeq2 v1.30.1 (76). Genes that were significantly altered between two conditions were defined by criteria of base mean > 100, log2 foldchange > 0.5 (or < −0.5) and padj < 0.05 by Wald test (DESeq2). Pathway analyses of genes were performed using Enrichr (79-81).

### ChRO-seq analyses

ChRO-seq reads were mapped to human genome hg38 by using an established pipeline (82). Briefly, PCR deduplication was performed by collapsing UMIs using PRINSEQ lite v0.20.2 (83). Adapters were trimmed from the 3’ end of remaining reads using Cutadapt v1.16 with a maximum 10% error rate, minimum 2 bp overlap, and minimum 20 quality score. Processed reads with a minimum length of 15 bp were mapped to the hg38 genome modified with the addition of a single copy of the human Pol I ribosomal RNA complete repeating unit (GenBank: U13369.1) using Burrows-Wheeler Aligner (BWA) v0.7.13 (84). The location of the RNA polymerase active site was represented by a single base that denotes the 5’ end of the nascent RNA, which corresponds to the position on the 3’ end of each sequenced read. The ChRO-seq reads were normalized by the length of gene bodies to transcripts per million (TPM). To visualize ChRO-seq signal, data were converted to bigwig format using bedtools and UCSC bedGraphToBigWig. Bigwig files within a sample category were then merged and normalized to a total signal of 1×10^6^ reads. Genomic loci snapshots were generated using R package Gviz (85).

### Quantification of gene transcription activity

Gene transcription activity was determined using ChRO-seq data. Gene body information was extract from GENCODE v33 (hg38). Genes with gene body smaller than 1 kb were excluded from this analysis. To quantify transcription activity of gene loci (gene length > 1kb), stranded ChRO-seq reads within gene coordinates were counted. Reads within 150 b downstream of transcription start site (TSS) were excluded to avoid bias of RNA polymerase pausing at the promoters (**Supplementary Figure 1**). Since mature miRNA species are transcribed from longer primary miRNAs transcripts, we used +/-5 kb flanking region of mature miRNA coordinate to determine transcription activity of miRNA species. Reads from transcriptional regulatory elements (TREs) that are present within gene bodies were also excluded to avoid bias of intragenic TRE activity. The TREs that are present in specific stage of directed differentiation experiments were defined previously in Hung et al. (26) using dREG (82). BEDTools was used to remove reads mapping to these TREs within gene bodies. The remaining reads were quantified using the R package bigwig (https://github.com/andrelmartins/bigWig). In differential transcription analysis, ChRO-seq raw counts of gene loci were analyzed through DESeq2 v1.30.1 (76).

### Define post-transcriptionally regulated genes

The development of this analysis pipeline was modified from (86). To define post-transcriptional regulation of genes, ChRO-seq and RNA-seq were analyzed through the two-factor analysis of DESeq2 v1.30.1 (76). Specifically, in the two-factor analysis, one factor referred to cell identities during a stage transition (e.g., duodenal spheroids versus definitive endoderm) and the other factor referred to type of assays (RNA-seq versus ChRO-seq). In this full model that contains these two factors and the interaction term between the two, the interaction term of the DESeq2 result was extracted to query the differences between assay types (RNA-seq, ChRO-seq) during a stage transition. Genes that were subject to significant post-transcriptionally regulation (become post-transcriptionally stable or unstable during a stage transition) were defined by criteria of log2 foldchange > 0.5 (or < −0.5) and padj < 0.05 in the interaction term by Wald test (DESeq2). To identify genes that had little changes at the transcriptional level (ChRO-seq) but significant changes at the steady state level (RNA-seq) during a stage transition, the post-transcriptionally regulated genes were filtered with additional criteria of padj > 0.1 in ChRO-seq differential expression analysis, padj < 0.05 in RNA-seq differential expression analysis, and log2 foldchange > 0.5 (or < −0.5) in RNA-seq differential expression analysis. In this study, only genes with ChRO-seq TPM > 20 and RNA-seq base mean > 100 were presented.

### miRhub analysis

The bioinformatic tool miRhub (87) was used to identify candidate master miRNA regulators of genes that were subject to post-transcriptional regulation in this study. miRhub uses the TargetScan algorithm to predict target sites for miRNAs of interest in the 3′ untranslated regions (UTRs) of a given gene set and then determines by Monte Carlo simulation for each miRNA whether the number and strength of predicted targets is significantly greater than expected by chance. The Monte Carlo simulation in a miRhub analysis was performed with 1000 iterations, with a set of randomly selected human genes each time, to generate a background distribution of the predicted targeting scores for each miRNA. These score distributions were then used to calculate an empirical p-value of the targeting score for each miRNA in a given gene set. In the present study, miRhub analyses were performed with predicated miRNA target sites that were present in humans and also preserved in one additional species (chicken, dog, mouse, rat)(using miRhub cons1 mode). Predicted miRNA target sites (species and sequence conservation) were obtained from TargetScan (88).

### Single cell RNA-seq analysis

Single cell sequencing reads from age and sex matched wildtype and miR-375 knockout mice (12-month-old; male) were aligned to the mouse genome (10x genomics pre-built reference version mm10-1.2.0) and quantified using Cell Ranger (v3.0.1) to obtain cell by gene count matrices. Ambient RNA removal was carried out using SoupX (v1.5.2). Subsequent filtering, data integration, clustering, and visualization of the data was completed using Seurat (v4.0.5). To maintain data quality, cells with less than 1500 detected genes or greater than 25% of the reads mapping to mitochondrial genes were discarded. 702 potential doublets, which were detected using scDblFinder (v1.4.0), were discarded, resulting in 11697 singlets for downstream analyses. To control sequencing depth, counts were normalized using sctransform (89) and glmGamPoi (v1.2.0). Samples were merged using the Seurat integration anchor workflow based on the 3000 most variable genes. Clustering and identification of nearest neighbors relied on 30 principal component analysis (PCA) dimensions. Cells were clustered at a resolution of 0.7, resulting in 17 cell clusters in the dataset. Clustering visualization was performed using uniform manifold approximation and projection (UMAP). Highly enriched markers for each cluster were determined using the Seurat function FindAllMarkers (log fold change > 0.5; using MAST method (90)), which compares the gene expression within a cluster with all other clusters. Identity of each of the cluster was assigned using addModuleScore function comparing highly enriched markers with known markers of intestinal epithelial cell types (curated from Habor et al., (91)) and classical immune cell markers (specifically *Gzma, Gzmb, Itgae, Ccl5, Cd7, Cd69, Cd3g* and Cd8a). Differential expression of genes between wildtype and miR-375 knockout samples was determined using function FindMarkers (using MAST method).

### Statistics

Statistical analyses were performed using R (4.0.4). In sequencing studies, statistical significance was determined using DESeq2, where P values were calculated by Wald test and adjusted using Benjamin and Hochberg (BH) method. Statistical significance of quantitative PCR was determined using unpaired two sample t-test unless otherwise noted. The P values in pathway enrichment analysis by Enrichr were calculated using Fisher’s exact test. Values of p < 0.05 (or adjusted p < 0.05 if in sequencing studies) were considered statistically significant. ∗p < 0.05, ∗∗p < 0.01, ∗∗∗p < 0.001.

## Data availability

Raw and processed data generated in the sequencing studies will be made available upon publication.

## Author contributions

Conceptualization and investigation, Y.-H.H., and P.S.; Formal analyses and bioinformatics, Y.-H.H.; Experiments, Y.-H.H., M.C., S.H., M.T.S. and J.V.; Data curation, Y-H.H., M.K., M.T.S., and J.V.; Mouse colony, R.L.C. and V.D.R.; Supporting and Resources, J.C.S.; Writing (original draft), Y.-H.H. and P.S.; Review and editing, Y.-H.H., M.C., J.R.S. and P.S.; Supervision, J.R.S. and P.S.

## Acknowledgements

We thank Dr. Zhao Li and the Greehey Children’s Cancer Research Institute at University of Texas Health Science Center at San Antonio for small RNA library preparation and sequencing; Dr. Peter Schweitzer and the Genomics Facility at Cornell University.

## Figure legends

**Supplementary Figure 1.**
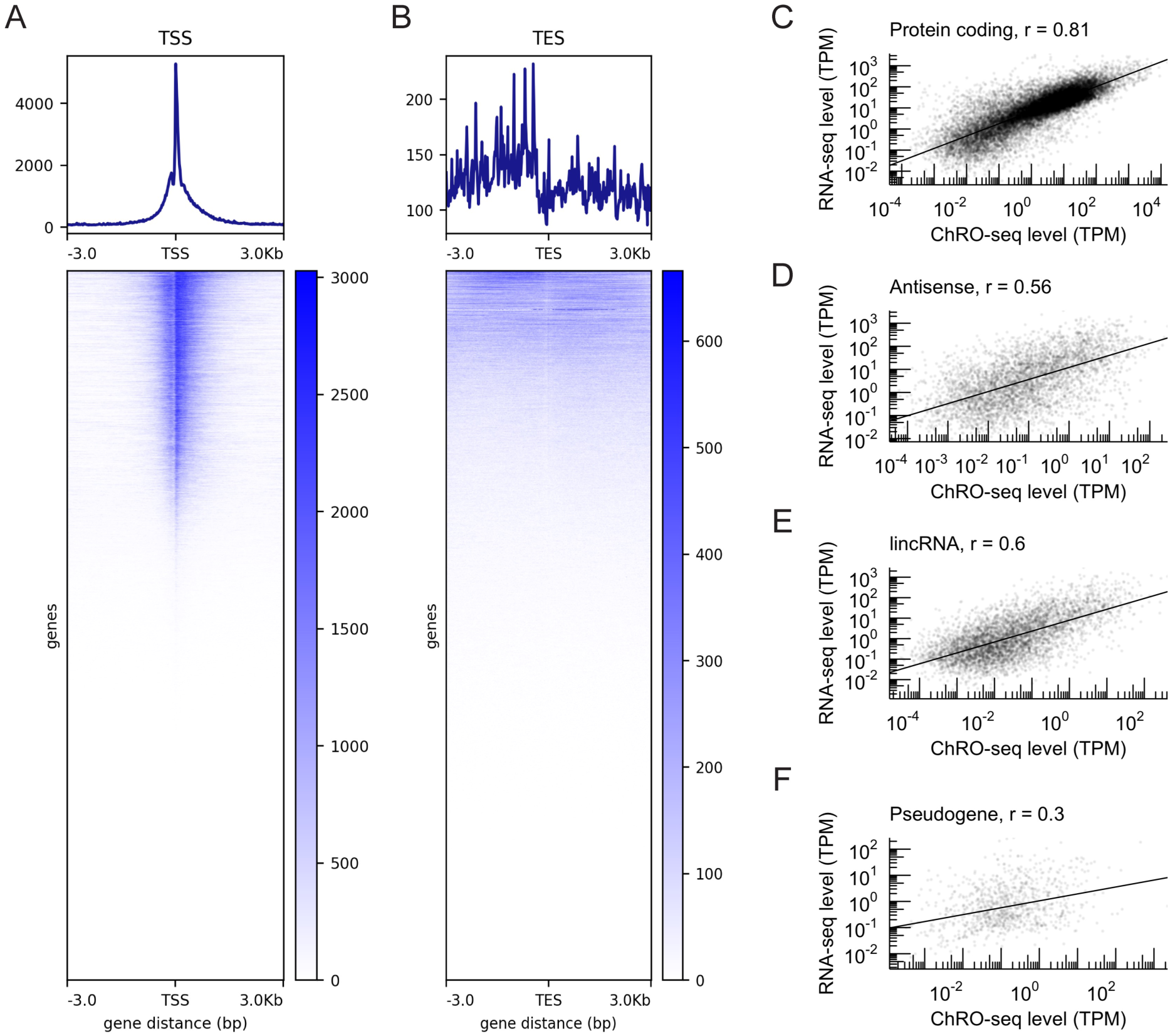
Correlation analysis between ChRO-seq and RNA-seq in the hPSC-HIO model. (A) ChRO-seq intensity around transcription start sites (TSSs). (B) ChRO-seq intensity around transcription termination sites (TESs). ChRO-seq signal per stage was normalized by total bigwig signal of 10^7^ before merging all four stages for this analysis. (C-F) Spearman correlation between ChRO-seq (TPM) and RNA-seq (TPM) across different gene classes, including protein coding (C), antisense (D), lncRNA (E) and pseudogene (F). In (C-F), ChRO-seq reads within 150 bp window of TSS downstream was excluded for TPM calculation and genes of which gene body is shorter than 1 kb were excluded. ChRO-seq study: hESC, n = 3; DE, n = 4; Duo spheroid (Duo), n = 3; Ile spheroid (Ile), n = 3. RNA-seq study: hESC, n = 2; DE, n = 3; Duo, n = 6; Ile, n = 4.

**Supplementary Figure 2.**
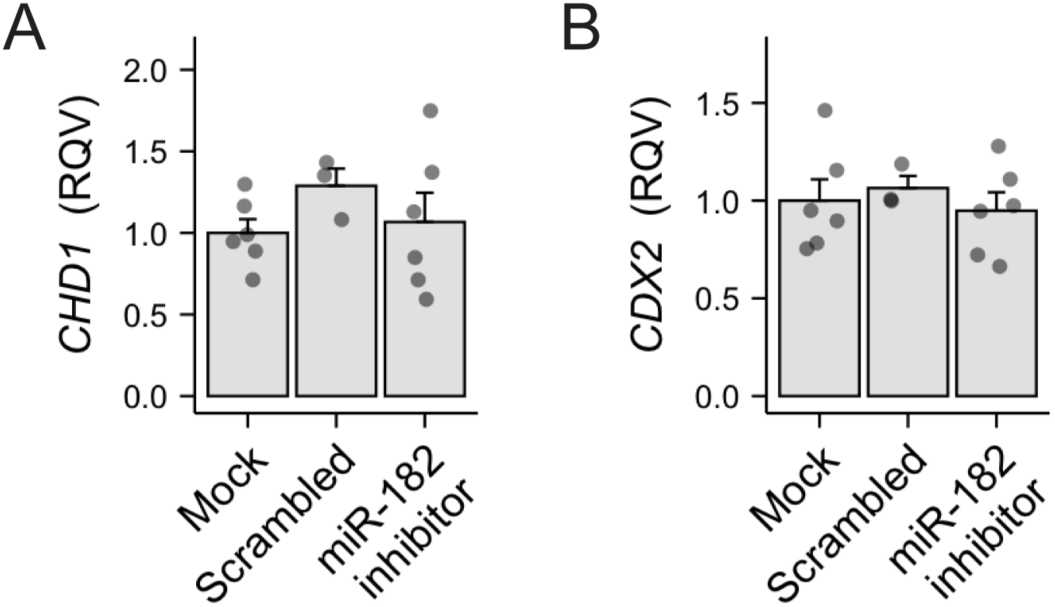
miRhub analyses identified candidate miRNA regulators in the hPSC-HIO model. (A-C) MiRhub analyses assessing the impacts of depleted miRNAs on the stable genes (left panel) and the impacts of enriched miRNAs on the unstable genes (left panel) during the event of DE formation (A), SI lineage formation (B) and ileal specification (C). Dash line denotes P-value = 0.05 in miRhub analyses.

**Supplementary Figure 3.**
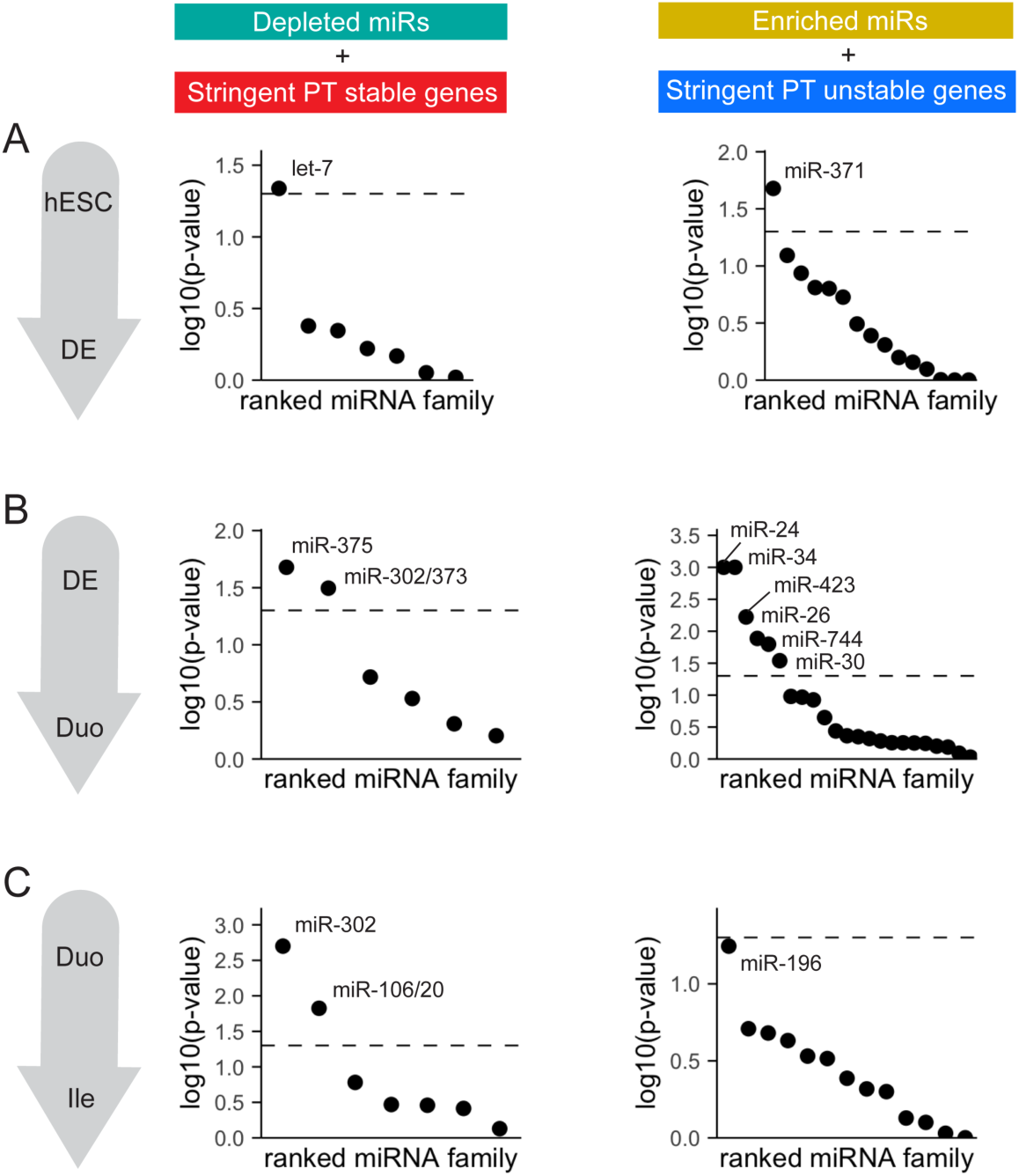
scRNA-seq analyses of the intestinal epithelial cells from adult WT and 375KO mice. (A) UMAP of initial clustering analysis with intestinal epithelial cells of adult WT and whole body miR-375 knockout (375KO) mice. (B) UMAP overlain with % mitochondrial genes in the dataset. (C) UMAP overlain with genotype information (WT vs. 375KO) in the dataset. (D) UMAP overlain with cell cycle phase in the dataset. (E) Differential abundance test (using MiloR) showing similar cell type composition between WT and 375KO mice in the dataset. (F) Numbers of differential expressed genes between WT and 375KO for each cell cluster in the dataset.

**Supplementary Figure 4.**
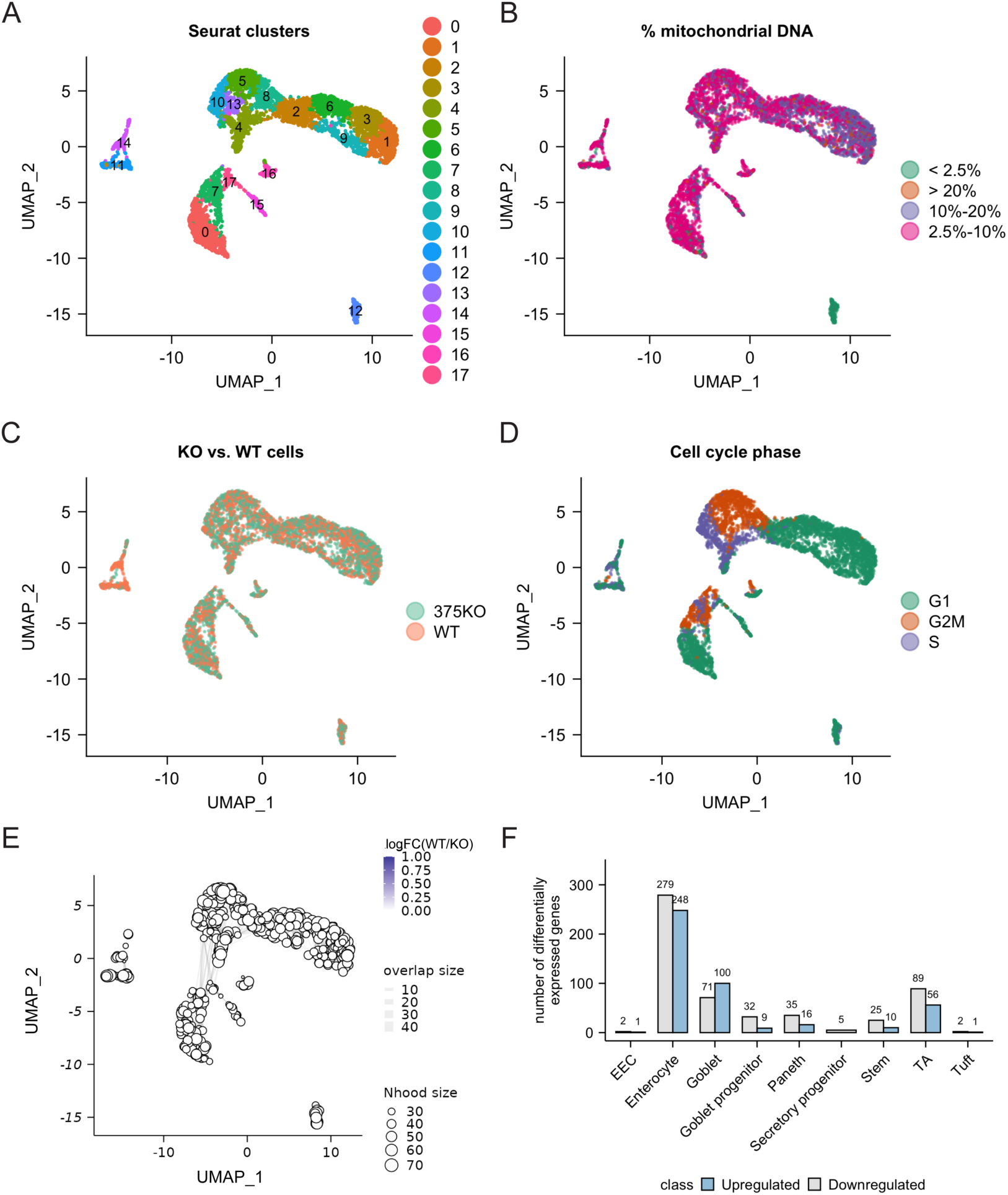
Low expression of *Arl4c* in the intestinal epithelial cells from adult WT and 375KO mice. (A) UMAP overlain of *Arl4c* expression in the intestinal epithelial cells of adult WT and whole body miR-375 knockout (375KO) mice. (B) Pseudo-bulk analysis showing a modest elevation of Arl4c in 375KO (n = 2) compared to WT (n = 2) mice (1.42 fold increase, P = 0.37 by two-tailed t-test).

**Supplementary Figure 5.**
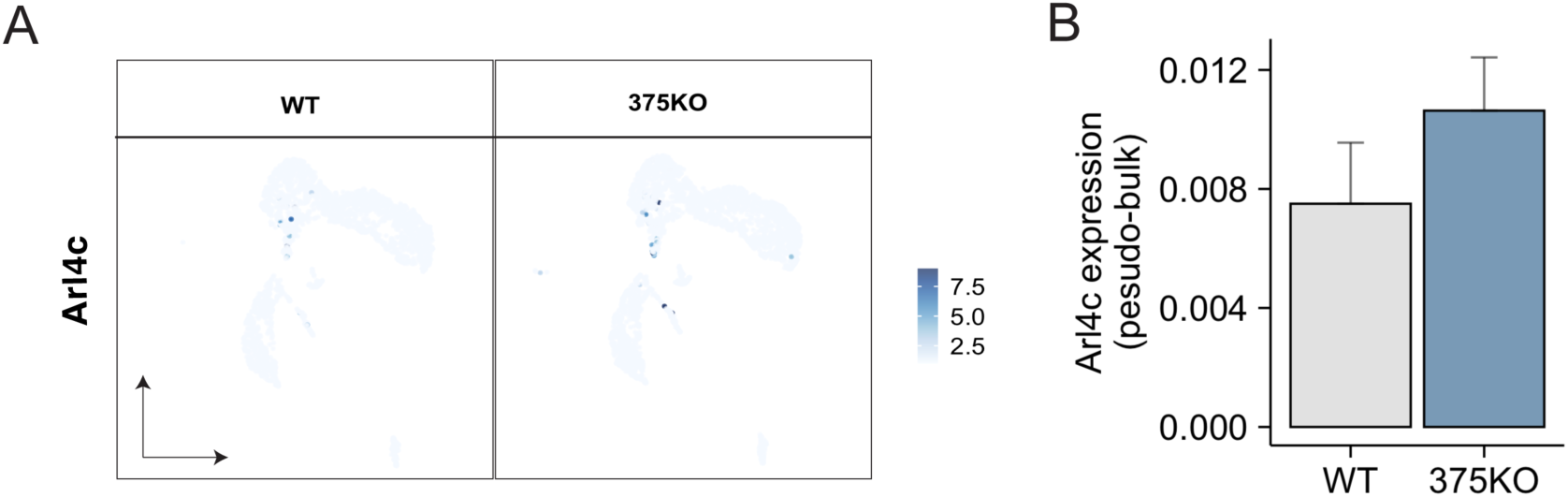

